# Epiblast formation by Tead-Yap-dependent expression of pluripotency factors and competitive elimination of unspecified cells

**DOI:** 10.1101/449397

**Authors:** Masakazu Hashimoto, Hiroshi Sasaki

## Abstract

The epiblast is a pluripotent cell population first formed in preimplantation embryos and its quality is important for proper development. Here, we examined the mechanisms of epiblast formation and found that the Hippo pathway transcription factor Tead and its coactivator Yap regulate expression of pluripotency factors. After specification of the inner cell mass, Yap accumulates in the nuclei and activates Tead. Tead activity is required for strong expression of pluripotency factors and is variable in the forming epiblast. Cells showing low Tead activity are eliminated from the epiblast through cell competition. Pluripotency factor expression and Myc control cell competition downstream of Tead activity. Cell competition eliminates unspecified cells and is required for proper organization of the epiblast. These results suggest that induction of pluripotency factors by Tead activity and elimination of unspecified cells via cell competition ensure the production of an epiblast with naïve pluripotency.

## Introduction

During preimplantation development, mouse embryos form a cyst-like structure known as the blastocyst. By the early blastocyst stage, embryos have formed two types of cells: the trophectoderm (TE) and inner cell mass (ICM). The ICM further differentiates into the epiblast and primitive endoderm by the late blastocyst stage. The TE and primitive endoderm later form extraembryonic tissues that support embryonic development. The epiblast at this stage has a naïve pluripotency and later forms all embryonic tissues and germ cells. Because a small number (approximately 10) of naïve epiblast cells produce the entire body, it is important that all of these cells are of high quality. Recent studies revealed that cell fate specification to the epiblast and primitive endoderm is a dynamic process involving heterogeneities and fluctuations in gene expression, elimination of cells via apoptosis, and asynchronous cell fate specification (Ohnishi et al., 2014; Plusa et al., 2008; Saiz et al., 2016; Xenopoulos et al., 2015). How this dynamic process leads to the production of high-quality epiblast cells is an important question to be addressed.

During development, developmental fluctuations and/or developmental errors can lead to the production of abnormal cells, which must be eliminated from the tissues to maintain tissue integrity. One mechanism by which this elimination occurs is known as cell competition (Claveria and Torres, 2016; Di Gregorio et al., 2016; Merino et al., 2016). Cell competition is a short-range intercellular communication in which cells compare their fitness with that of neighboring cells, and those with relatively lower fitness are eliminated. Pioneering studies in *Drosophila* imaginal discs revealed the factors and signaling pathways controlling cell competition; among them, the transcription factor Myc and Hippo signaling pathway play important roles in regulating cell competition (de la Cova et al., 2004; Moreno and Basler, 2004; Neto-Silva et al., 2010; Ziosi et al., 2010). In mouse embryos, differential expression of Myc triggers cell competition in the postimplantation epiblast at approximately embryonic day 6.0 (E6.0), which eliminates cells that are prematurely primed for differentiation and/or cells with lower fitness (Claveria et al., 2013; Diaz-Diaz et al., 2017; Sancho et al., 2013).

In contrast, the roles of Hippo signaling in cell competition in mouse embryos remain unknown. Activation of the Hippo signaling pathway suppresses nuclear localization of the coactivator proteins Yap (encoded by *Yap1*) and Taz (encoded by *Wwtr1*) to inactivate Tead family transcription factors (Lei et al., 2008; Ota and Sasaki, 2008; Zhao et al., 2007). Mouse has four *Tead* genes (*Tead1–Tead4*), and the activity of Tead transcription factors (hereafter Tead activity) controls cell proliferation (Ota and Sasaki, 2008; Zhao et al., 2008). We previously showed that in NIH3T3 mouse embryonic fibroblasts, co-culture of the cells with different Tead activities triggers cell competition, in which the cells with higher Tead activity were retained (Mamada et al., 2015). Tead controls Myc expression, and Tead activity and Myc cooperatively regulate cell competition. However, the effects of Hippo signaling on cell competition vary in different experimental system. Knock-down of YAP1 did not cause cell competition in U2OS human osteosarcoma cells (Penzo-Mendez et al., 2015), while cells expressing active YAP1 were eliminated through apical extrusion in Madin-Darby canine kidney cells (Chiba et al., 2016).

Another important question regarding epiblast formation is how the cells acquire naïve pluripotency in embryos. Studies of embryonic stem (ES) cells revealed the molecular mechanisms regulating the maintenance of naïve pluripotency (Martello and Smith, 2014; Nichols and Smith, 2009; Niwa, 2007, 2018; Smith, 2017). Some of these mechanisms are also used in formation of the naïve epiblast. However, the roles of Hippo signaling pathway in the formation and maintenance of naïve pluripotency remain unclear. While some studies showed that Tead-Yap supports pluripotency in the self-renewal of ES cells and induction of induced pluripotent stem cells (Li et al., 2013; Lian et al., 2010; Tamm et al., 2011), other reports showed that these proteins are dispensable for ES cell self-renewal (Azzolin et al., 2014; Chung et al., 2016). In pre-implantation embryos, Hippo signaling regulates the first cell fate specification to the TE and ICM (Sasaki, 2017), but no studies have examined the role of Hippo signaling during formation of the epiblast from the ICM. Thus, it remains unclear whether Hippo signaling and/or Tead-Yap are involved in the acquisition of naïve pluripotency during epiblast formation.

In this study, we show that differences in Tead activities trigger cell competition in the forming epiblast in which cells with lower Tead activity were eliminated. In normal embryos, attenuation of Hippo signaling increased Tead activity in the forming epiblast; this activity is required for high-level expression of pluripotency factors. Cell competition eliminates unspecified and low pluripotency factor cells produced by variations in Tead activities. These results suggest that Tead activity promotes pluripotency, and cell competition eliminates unspecified cells to establish an epiblast with naïve pluripotency.

## Results

### Cell competition eliminates *Tead1* mutant cells from epiblast by E6.5

To examine the possible involvement of Tead activity in cell competition during mouse development, we produced mosaic embryos consisting of wild-type cells and *Tead1* mutant cells (wild-type ⇔ *Tead1*^−/−^ embryos). We selected *Tead1*^−/−^ because NIH3T3 cells with *Tead1* knockdown showed low Tead activity and were eliminated during co-culture with normal cells (Mamada et al., 2015). To generate mosaic embryos, we injected CRISPR/Cas9 protein, two adjacent *Tead1* sgRNAs, and *Cre* mRNA into one blastomere of two-cell stage embryos of the ROSA26 knock-in Cre reporter mouse R26GRR (Hasegawa et al., 2013). This manipulation produced mosaic embryos consisting of wild-type cells labeled with green fluorescence and *Tead1*^−/−^ cells labeled with red fluorescence (Figure 1A). The efficiencies of *Tead1* knock-out and color conversion by Cre recombinase were 100% (n = 18/18) and 96% (n = 78/81), respectively (Figure S1A-D). At E6.5, *Tead1*^−/−^ cells failed to contribute to the epiblast (Figure 1C, D), but contributed to the visceral endoderm and extraembryonic ectoderm, which are extraembryonic tissues (Figures 1C and S1E, F). Because control (wild-type ⇔ wild-type) mosaic embryos showed mixing of the two colors throughout conceptus (Figure 1B, D), a biased distribution was created by *Tead1* mutation. The absence of *Tead1*^−/−^ cells in the epiblast is not a cell autonomous defect, as simple *Tead1*^−/−^ embryos survive up to E11.5 (Chen et al., 1994; Sawada et al., 2008). We also confirmed that the E6.5 *Tead1*^−/−^ embryos produced by genome editing at the one-cell stage showed a normal morphology and developmental rate (Figures 1E, F and S1G, H). These results suggest *Tead1*^−/−^ cells are absent from the epiblast because of cell competition between wild-type and *Tead1*^−/−^ cells.

**Figure 1.**
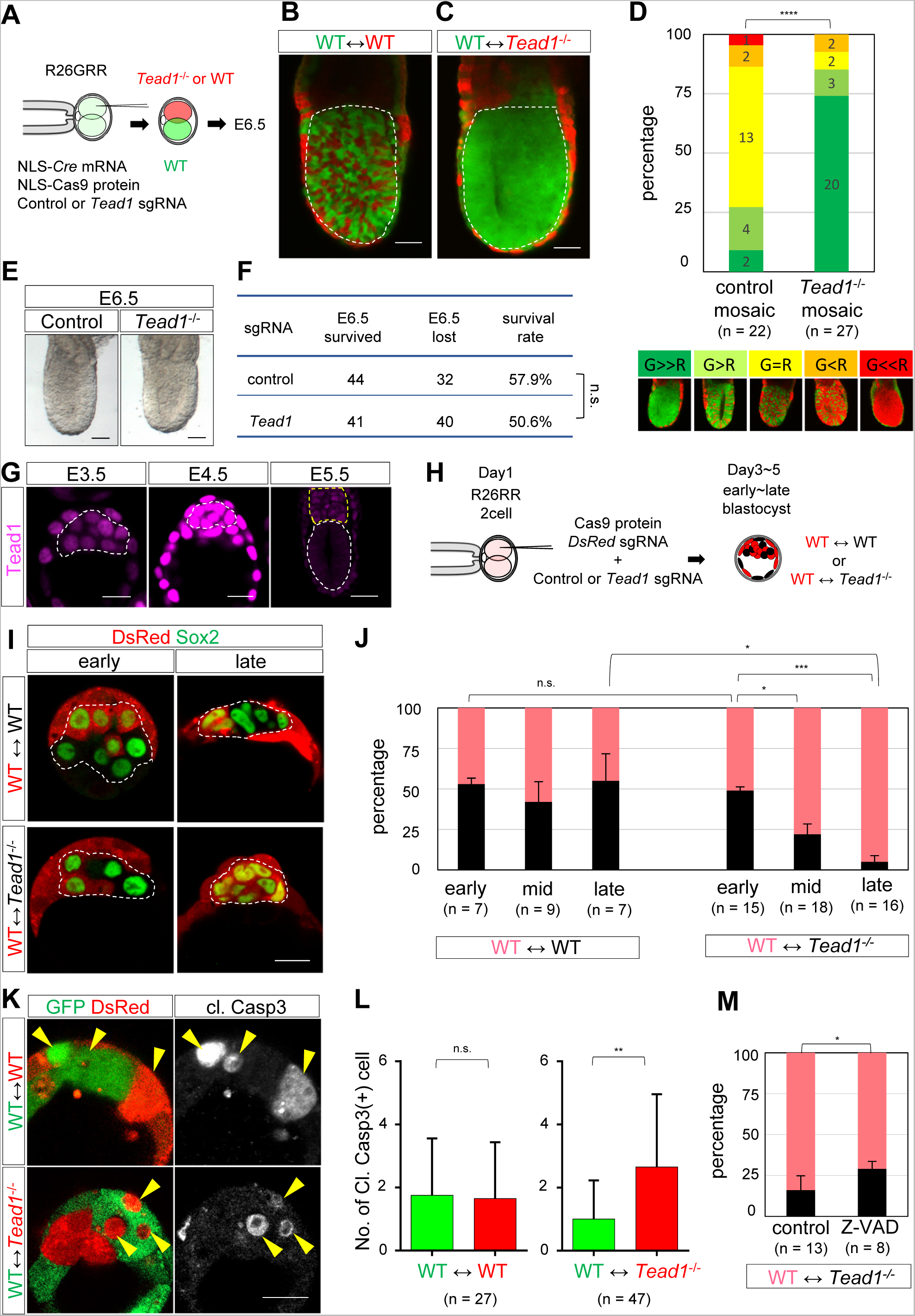
*Tead1*^−/−^ cells are eliminated from epiblast via cell competition. (A-F) Elimination *of Tead1^−/−^* cells from epiblast in wild-type ⇔ *Tead1*^−/−^ embryos at E6.5. (A) Scheme of experiments. (B, C) Wild-type (WT) ⇔ WT control (B) and WT ⇔ *Tead1*^−/−^ (C) embryos at E6.5. Dashed lines mark epiblasts. (D) (top) Distribution of two cell types in epiblasts. (bottom) Representative embryo of each category showing distribution of two cell types in epiblast. (E) Morphology of control and *Tead1*^−/−^ embryos at E6.5. (F) Developmental rate *of Tead1^−/−^* embryos at E6.5. (G) Distribution of Tead1 proteins in embryos. Dashed lines indicate ICM (E3.5), epiblasts (E4.5 and E5.5), and extra-embryonic ectoderm (E5.5). (H) Scheme of experiments. (I) Distribution of WT and *Tead1*^−/−^ cells in ICM/epiblast. Dashed lines indicate ICM/epiblast. (J) Quantification *of Tead1^−/−^* cells in ICM/epiblast. (K) Distribution of apoptotic cells in mosaic embryos. Arrows indicate apoptotic cells. (L) Quantification of apoptotic cells. (M) Z-VAD-treatment increased contribution *of Tead1^−/−^* cells in epiblast. Data are represented as mean ± SEM (J, M) or SD (L). Chi-square test (D, F), One-way ANOVA followed by Dunn’s multiple comparison test (J), and Student’s *t*-test (L, M). *\* p* < 0.05, *\*\* p <* 0.01, *\*\*\* p <* 0.001, *\*\*\*\* p <* 0.0001. Scale bars are 50 µm for panels B, C, E, G (E5.5) and 20 µm for G (E3.5, E4.5), I, K. See also Figure S1.

### Elimination of *Tead1*^−/−^ cells occurs during epiblast formation

We next investigated the timing of cell competition. Because *Tead1* mRNA is not expressed in the epiblast of E6.5 embryos (Sawada et al., 2005), we evaluated the distribution of Tead1 proteins in earlier stage embryos. Tead1 proteins were widely present in preimplantation embryos with strong expression at E4.5, while no Tead1 signal was observed in the epiblast of early post-implantation embryos at E5.5 (Figure 1G). These results suggest that cell competition takes place in preimplantation embryos. In subsequent experiments, unless otherwise specified, we used transgenic mice expressing red fluorescent protein (R26RR) and co-injected sgRNAs for *Tead1* and *DsRed to* remove DsRed signals in *Tead1*^−/−^ cells (Figure 1H). The resulting mosaic embryos consisted of red wild-type cells and non-fluorescent (black) *Tead1*^−/−^ cells (Figure 1H). The knockout efficiency of *DsRed* was 100% (n = 18/18) (Figure S1A,B). We used the pluripotency factor Sox2 as a marker of the ICM and epiblast because of its specific expression in these cells in early and late blastocysts, respectively. At the early blastocyst (32-63-cell) stage, the ICM of the wild-type ⇔ *Tead1*^−/−^ embryos consisted of similar numbers of wild-type and the *Tead1 ^−/−^* cells (Figure 1I, J). The ratio of *Tead1*^−/−^ cells gradually decreased from the mid blastocyst (64-127-cell) stage and at the late blastocyst (>128-cell) stage, with a contribution to the epiblast cells of 5%. Therefore, elimination occurs between the early and late blastocyst stages (Figure 1I, J).

To clarify the process of mutant cell elimination, we examined the involvement of apoptosis. We used R26GRR mice for this experiment (Figure 1A). At the mid-blastocyst stage, control embryos showed similar numbers of cleaved caspase 3-positive apoptotic cells in both injected and uninjected epiblast cells, while wild-type ⇔ *Tead1*^−/−^ embryos contained larger numbers of apoptotic cells in the *Tead1*^−/−^ epiblast cells (Figure 1K, L). Treatment of mosaic embryos with a pan-caspase inhibitor (Z-VAD-FMK, hereafter Z-VAD) from the early blastocyst stage increased the ratio *of Tead1^−/−^* cells in the epiblast of late blastocyst stage embryos (Figure 1M). Taken together, these results suggest that *Tead1*-dependent cell competition occurs during epiblast formation between the early and late blastocyst stages, and that elimination of *Tead1*^−/−^ cells involves caspase-dependent apoptosis.

### Tead becomes active in the forming epiblast

Elimination of *Tead1*^−/−^ cells suggests that differences in Tead activities caused cell competition in the forming epiblast cells. However, we previously showed that in the ICM of early blastocyst stage embryos, active Hippo signaling inactivated Tead proteins by excluding Yap from the nuclei (Hirate et al., 2012; Nishioka et al., 2009). To clarify whether Tead proteins are also inactive in the forming epiblast, we first examined the expression of Yap during the blastocyst stages. Confirming our previous observations, Yap was mostly present in the cytoplasm in the ICM of early blastocysts. At the mid-blastocyst stage, however, Yap began to accumulate in the nuclei and at the late blastocyst stage, Sox2-positive epiblast cells showed strong nuclear Yap signals, suggesting that Tead activity gradually increased during the formation of epiblast cells (Figure 2A, A’, B). In support of activation of Tead proteins, active Yap proteins, which are non-phosphorylated and localize to the nuclei (Zhao et al., 2007), were present in the epiblast of late blastocysts (Figure 2C, D). Furthermore, Amotl2, a target gene of Tead-Yap in ES cells (Papaspyropoulos et al., 2018), was expressed in the epiblast (Figure 2E, F). Thus, Tead is activated during epiblast formation.

**Figure 2.**
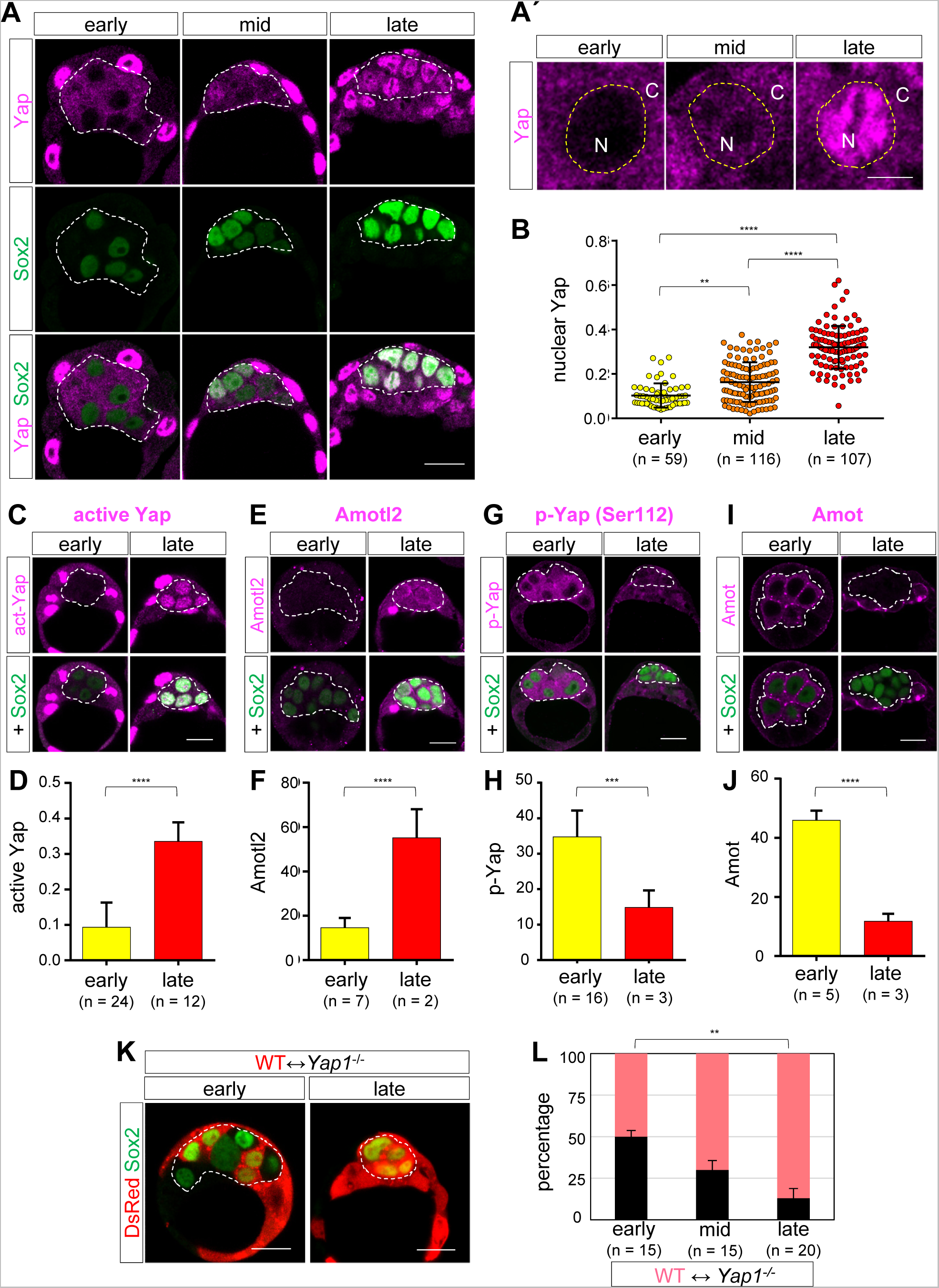
Attenuation of Hippo signaling promotes nuclear accumulation of Yap in epiblast. (A) Normal embryos showing gradual increase of nuclear Yap signals in forming epiblast. Dashed lines indicate Sox2-positive ICM/epiblast cells. (A’) Enlargement of (A). (B) Quantification of nuclear Yap signals. (C-J) Changes in Hippo signal-related proteins between ICM and epiblast. Representative embryos and quantification of active Yap (C, D), Amotl2 (E, F), phospho-S112-Yap (G, H), and Amot (I, J). Dashed lines indicate Sox2-positive ICM/epiblast. (K) WT ⇔ *Yap1^−/−^* embryos showing gradual elimination of *Yap1^−/−^* cells from forming epiblast. (L) Quantification of the percentage of *Yap1^−/−^* cells. Data are represented as the mean ± SD (B, D, F, H, J) or SEM (L). One-way ANOVA followed by Dunn’s multiple comparison test (B, L), Student’s *t*-test (D, F, H, J). \*\**p* < 0.01, *** *p* < 0.001, **** *p* < 0.0001 Scale bars are 20 µm for panels A, C, E, G, I, K and 5 µm for A’. See also Figure S2.

The subcellular distribution of Yap is regulated by both Hippo signal-dependent and independent mechanisms, e. g. mechanical tension (Dupont et al., 2011; Elosegui-Artola et al., 2017). Activation of the Hippo signaling pathway results in the phosphorylation of serine 112 of Yap (p-Yap) by Lats1/2 kinase (Zhao et al., 2007). A strong p-Yap signal was observed in the ICM of the early blastocysts, but the epiblast of late blastocysts clearly showed weaker signals, suggesting that reduced Hippo signaling promoted nuclear localization of Yap (Figure 2G, H). Junction-associated proteins, angiomotin (Amot) family proteins, are required for activation of the Hippo pathway in the ICM (Hirate et al., 2013). Amot is strongly expressed in the ICM of early blastocysts but is absent in the epiblast of late blastocysts (Figure 2I, J). This strong reduction in Amot likely contributes to the reduction of Hippo signaling. In summary, after specification of the ICM fate, Hippo signaling is attenuated and Tead activity gradually increases in the forming epiblast cells.

To further evaluate the importance of Tead activities in cell competition, we generated wild-type ⇔ *Yap1^−/−^* mosaic embryos (Figure S2A, B), in which *Yap1^−/−^* cells have lower Tead activity. As observed for wild-type ⇔ *Tead1*^−/−^ embryos, *Yap1^−/−^* cells were gradually eliminated from the forming epiblast after the mid-blastocyst stage (Figure 2K, L). Because *Yap1^−/−^* embryos develop up to E8.5 (Morin-Kensicki et al., 2006), elimination of *Yap1^−/−^* cells from the forming epiblast is a consequence of cell competition with wild-type cells. Taken together, during epiblast formation, Tead activity gradually increases, and differences in Tead activities induce cell competition to eliminate cells with lower Tead activities.

### Tead activity promotes pluripotency

Cells with low Tead activity are eliminated from the epiblast via cell competition. To understand the characteristics of the eliminated cells, we next examined the role of Tead activity in epiblast formation. In parallel with the increase in nuclear Yap, expression of Sox2 gradually increased during the blastocyst stages (Figures 2A, B and 3A). Quantification of the signal intensities revealed a strong correlation between nuclear Yap levels and Sox2 expression levels (Figure 3B). Weaker but significant correlations with nuclear Yap levels were also observed for the expression levels of other core pluripotency factors, Oct3/4 and Nanog (Figure 3C-F). Nanog expression during the early blastocyst stage was stochastic (Dietrich and Hiiragi, 2007; Plusa et al., 2008) and unrelated to nuclear Yap (Figure 3E, F boxed), while expression of Nanog at the mid and late blastocyst stages showed a stronger correlation with nuclear Yap (Figure 3F). Therefore, Tead activity is likely involved in the acquisition of pluripotency to form epiblast cells.

**Figure 3.**
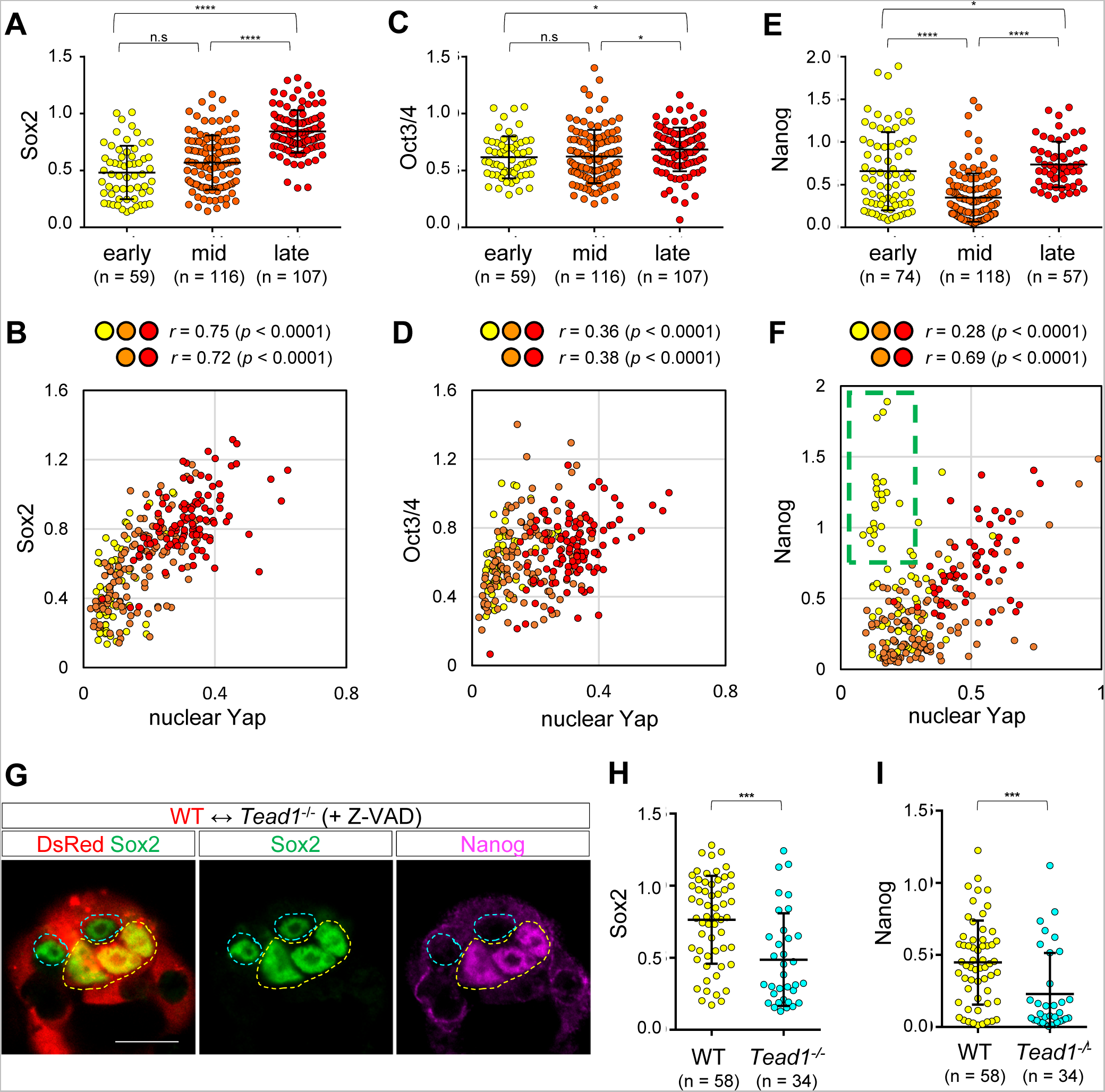
Tead activity promotes pluripotency factor expression in forming epiblast. (A-F) Correlations between signals of pluripotency factors and nuclear Yap. Signal quantification of pluripotency factors, Sox2 (A), Oct3/4 (C), and Nanog (E), in Sox2-positive ICM/epiblast cells. Relationships between signals of nuclear Yap and pluripotency factors, Sox2 (B), Oct3/4 (D), and Nanog (F). Data from early, mid, and late blastocyst stages are labeled with yellow, orange, and red colors, respectively. Correlation coefficients and *p* values are shown above the graphs. (G-I) Tead activity is required for strong expression of pluripotency factors. (G) Z-VAD-treated WT ⇔ *Tead1*^−/−^ embryo showing weaker expression of pluripotency factors, Sox2 and Nanog, in *Tead1*^−/−^ cells. Yellow and cyan dashed lines indicate wild-type and *Tead1*^−/−^ cells, respectively. Scale bar represents 20 µm. Quantification of Sox2 (H) and Nanog (I) signals in WT and *Tead1*^−/−^ cells at late blastocyst stage. Data are represented as the mean ± SD. One-way ANOVA followed by Dunn’s multiple comparison test (A, C, E), Mann-Whitney *U* test (H, I). \**p* < 0.05, *** *p* < 0.001, **** *p* < 0.0001.

To test this hypothesis, we treated wild-type ⇔ *Tead1*^−/−^ embryos with Z-VAD from the early blastocyst stage to suppress elimination *of Tead1^−/−^* cells via cell competition. Z-VAD-treatment did not completely suppress elimination, but the remaining *Tead1*^−/−^ cells showed significantly weaker expression of Sox2 and Nanog compared to wild-type cells at the late-blastocyst stage (Figure 3G-I), suggesting that Tead activity is required for strong expression of pluripotency factors in epiblast cells. This also indicates that cell competition eliminates cells with low pluripotency factor expression.

### Tead activity regulates endogenous cell competition in epiblast

Our results revealed that cell competition was induced in the forming epiblast when cells with low Tead activity were experimentally introduced. We next examined whether similar cell competition occurs in normal embryos. Consistent with a previous observation (Copp, 1978), most of the ICM of freshly recovered blastocysts contained cleaved caspase 3-positive apoptotic cells (Figure 4A, B). A particularly high frequency of apoptosis was observed in early mid-blastocyst stage (64–95-cell) embryos (Figure 4B). At the mid-blastocyst stage, nuclear Yap levels and the expression of pluripotency factors in the ICM cells were highly variable among ICM cells in an embryo, and 70% of embryos showed a significant correlation between nuclear Yap levels and Sox2 expression (Figure 4C, D, Figure S3A–F). Therefore, sharp differences in Tead activities/pluripotency factors likely trigger cell competition. In support of this hypothesis, at the mid-blastocyst stage, the cleaved caspase3-positive ICM cells showed weak nuclear Yap and Sox2 signals (Figure 4E–G). At the late blastocyst stage, the expression of Sox2 and Oct3/4 in the epiblast was stronger (Figure 3A, C) and variations were smaller (Figure 4H, I) than those at the mid blastocyst stage, respectively. These results are consistent with the hypothesis that at the mid blastocyst stage, variations in Tead activities cause variations in pluripotency factor expression and trigger cell competition, eliminating cells with low Tead activity/low pluripotency factors from the forming epiblast.

**Figure 4.**
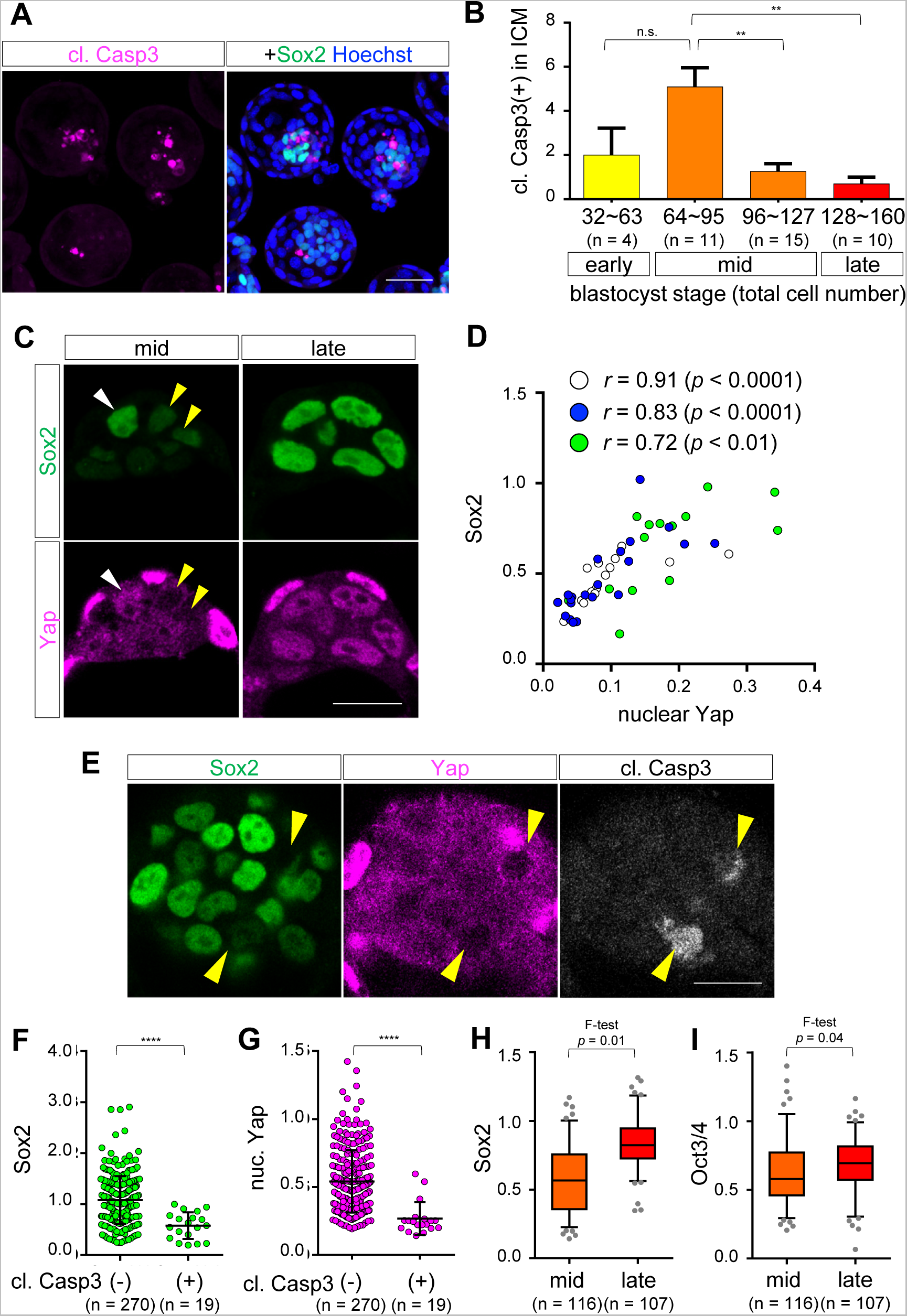
Endogenous cell competition triggered by Tead activity takes place at mid blastocyst stage. (A) Wild-type embryos showing cleaved caspase 3 (cl. Casp3) signals in ICM/epiblast. (B) Quantification of apoptotic cells in the ICM/epiblast. (C) Variation in nuclear Yap and Sox2 signals within an embryo. White and yellow arrowheads indicate the cells with strong and weak nuclear Yap/Sox2 signals, respectively. At the late blastocyst stage, the signals are less variable. (D) Quantification of nuclear Yap and Sox2 signals showing strong variation in the signal intensities within individual embryos at mid-blastocyst stage. The data derived from different embryos are labeled with different colors. (E) Cleaved caspase 3-positive cells showing weak nuclear Yap/Sox2 signals (yellow arrowheads). (F, G) Quantification of Sox2 (F) and nuclear Yap (G) signals. (H, I) Comparison of variations of Sox2 (H) and Oct3/4 (I) signals between mid and late blastocyst stages. Data are represented as the mean ± SD (B, F, G) or box and whisker plots (H, I). Box and the horizontal line indicate the range of central 50% and median, respectively. Whiskers represent the range from 5% to 95%. One-way ANOVA followed by Dunn’s multiple comparison test (B), Student’s *t*-test (F, G), *F*-test (H, I). * *p* < 0.05, ** *p* < 0.01, **** *p* < 0.0001. Scale bars are 50 µm for panel A and 20 µm for C, E. See also Figure S3.

### Pluripotency and Myc control cell competition downstream of Tead activity

Because Tead activity regulates expression of pluripotency factors, it is not clear whether Tead activity itself controls cell competition or pluripotency factors regulate cell competition. To address this issue, we manipulated the expression of pluripotency factors by treating the embryos with two kinase inhibitors known as “2i”. The 2i inhibitors, PD0325901 and CHIR99021, inhibit mitogen-activated protein kinase kinase (Mek) and glycogen synthase kinase-3 (Gsk3), respectively (Ying et al., 2008). As previously reported, in the presence of 2i, expression of pluripotency factors, particularly Nanog, in the epiblast was strongly upregulated (Nichols et al., 2009) (Figure S4A). All ICM cells became part of the epiblast, while no Sox17-positive primitive endoderm was formed (not shown). When wild-type ⇔ *Tead1*^−/−^ embryos were cultured in the presence of 2i, *Tead1*^−/−^ cells expressed high levels of Oct3/4 which were comparable to those in wild-type cells and elimination of *Tead1*^−/−^ cells was strongly suppressed (Figure 5A, B). Treatment of wild-type embryos with 2i also suppressed endogenous cell death as shown by the strong reduction in cleaved caspase3-positive cells in the ICM (Figures 5C, S4B). The 2i treatment increased expression of pluripotency factors in epiblast cells independently of Tead activity, as shown by the increased Sox2 expression without an increase in the nuclear Yap level in the epiblast (Figure 5D-F). These results suggest that expression of pluripotency factors, or most likely pluripotency, is a critical factor regulating cell competition downstream of Tead activity. However, because 2i-treatment also has multiple effects other than increasing pluripotency factors, we examined whether other mechanisms are involved in regulating cell competition.

**Figure 5.**
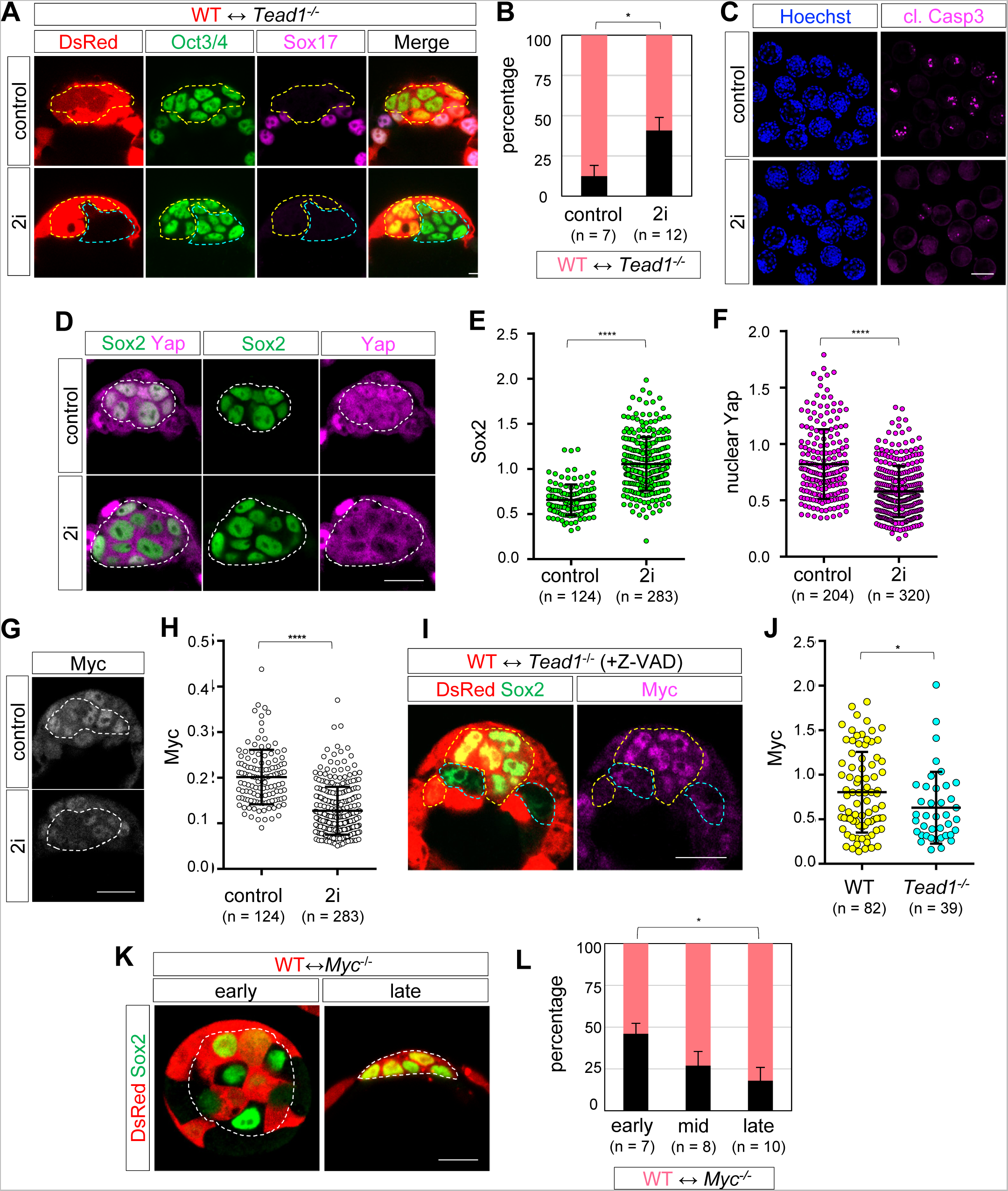
Pluripotency and Myc regulates cell competition downstream of Tead activity. (A) Distribution of wild-type (yellow dashed lines) and *Tead1*^−/−^ cells (cyan dashed lines) in the epiblast of wild-type ⇔ *Tead1*^−/−^ embryos. 2i-treatment suppressed elimination *of Tead1^−/−^* cells. (B) Quantification of percentages of *Tead1*^−/−^ cells in the epiblast. (C) Reduction of cleaved caspase 3-positive cells in 2i-treated wild-type embryos. (D) Effects of 2i-treatment on Sox2 expression and nuclear Yap. (E, F) Quantification of signal intensities of Sox2 (E) and nuclear Yap (F). (G) 2i-Treated wild-type embryo showing reduced Myc expression in ICM. (H) Quantification of Myc signals in the ICM. (I) Z-VAD-treated WT ⇔ *Tead1*^−/−^ embryo showing weaker expression of Myc in *Tead1*^−/−^ epiblast cells. (J) Quantification of Myc signals in epiblasts. (K) Changes in percentage *of Myc^−/−^* cells in the ICM/epiblast of wild-type ⇔ *Myc^−/−^* embryos. (L) Quantification of percentage of *Myc^−/−^* cells in ICM/epiblast. Data are represented as the mean ± SEM (B, L) or SD (E, F, H, J). Student’s *t*-test (B, E, F), Mann-Whitney *U* test (J), One-way ANOVA followed by Dunn’s multiple comparison test (L). * *p* < 0.05, **** *p* < 0.0001. Scale bars are 20 µm for panels A, D, G, I, K and 100 µm for C. See also Figures S4 and S5.

An alternative possibility is that the absence of primitive endoderm suppressed cell competition. To examine this possibility, we used *Gata6^−/−^* embryos, which lack primitive endoderm (Schrode et al., 2014). The *Gata6^−/−^* embryos freshly produced by genome editing at the one-cell stage did not form primitive endoderm (Figure S4C). In the *Gata6^−/−^* ⇔ *Tead1*^−/−^; *Gata6^−/−^* mosaic embryos, which were produced by *Gata6* knock-out at the one-cell stage followed by *Tead1* knock-out at the two-cell stage (Figure S4D), *Tead1*^−/−^;*Gata6^−/−^* cells were efficiently eliminated (Figure S4E, F). Therefore, the absence of primitive endoderm did not affect cell competition triggered by Tead activity.

Another possibility is that 2i treatment altered the expression of genes involved in cell competition. In blastocysts, Myc was expressed throughout the embryos and stronger and variable expression was observed in the epiblast, as reported previously (Claveria et al., 2013), and embryos cultured with 2i showed significantly weaker expression of Myc in the epiblast (Figure 5G, H). Because 2i treatment affected Myc expression, we next evaluated whether Myc is regulated by Tead activity. In wild-type ⇔ *Tead1*^−/−^ embryos treated with Z-VAD, *Tead1*^−/−^ cells showed weaker Myc expression (Figure 5I, J), suggesting that Tead activity controls Myc. Therefore, we examined whether differential Myc expression triggers cell competition. In wild-type ⇔ *Myc^−/−^* mosaic embryos, *Myc^−/−^* cells were gradually eliminated from the forming epiblast after the mid-blastocyst stage, resembling wild-type ⇔ *Tead1*^−/−^ embryos (Figures 5K, L, S5A, B). Because *Myc^−/−^* embryos survive up to E9.5 (Davis et al., 1993), cell competition with wild-type cells eliminated *Myc^−/−^* cells from the epiblast. These results suggest that Myc is involved in Tead activity-triggered cell competition.

Based on these results, we next examined whether inhibition of Myc suppresses Tead-activity triggered cell competition. Mice contain three *Myc* genes *(Myc, Mycn, Mycl).* To simultaneously inactivate all three Myc proteins, we treated wild-type ⇔ *Tead1*^−/−^ embryos with the pan-Myc inhibitor MYCi, which inhibits interactions between Myc family proteins and their common partner protein, Max. MYCi-treated embryos showed reduced expression of Myc, which was also observed in the absence of the Myc-Max interaction (Hishida et al., 2011) (Figure S5C, D). MYCi-treated wild-type ⇔ *Tead1*^−/−^ embryos failed to suppress elimination *of Tead1^−/−^* cells from the epiblast (Figure S5E, F), indicating that Myc is not the only factor regulating cell competition downstream of Tead activity. Therefore, it is likely that pluripotency and Myc cooperatively control cell competition downstream of Tead activity.

Because of the unavailability of techniques for manipulating pluripotency in the forming epiblast independently of Tead activity and/or Myc, we could not directly examine the role of pluripotency in Tead activity-triggered cell competition. Nevertheless, all experimental results described above are consistent with the hypothesis that pluripotency and Myc cooperatively control cell competition downstream of Tead activity.

### Cell competition eliminates unspecified and low-pluripotency cells

Cell competition eliminates cells with low levels of pluripotency factors, but it remains unclear whether these cells are epiblast cells or if these cells have an alternative fate of the primitive endoderm. Indeed, it was reported that primitive endoderm cells mislocated inside the epiblast undergo apoptosis (Plusa et al., 2008). To address this issue, we examined the expression of the epiblast marker Sox2 and primitive endoderm marker Sox17 at the late blastocyst stage. In wild-type ⇔ *Tead1 ^−/−^* embryos, the wild-type inner cells strongly expressed either Sox2 or Sox17, indicating that the cells were specified to either the epiblast or primitive endoderm (Figure 6A, B). Consistent with elimination of the *Tead1*^−/−^ cells, the number of *Tead1*^−/−^ cells was clearly smaller than that of wild-type cells, but the remaining cells were also single-positive for Sox2 or Sox17 (Figure 6C), indicating specification to the epiblast and primitive endoderm. To characterize the identities of the eliminated cells, we next treated the wild-type ⇔ *Tead1*^−/−^ embryos with Z-VAD from the early blastocyst stage to partially suppress apoptosis (Figure 6D-F). In *Tead1 ^−/−^* cells, in addition to the normally specified Sox2 or Sox17 single-positive cells, a new population of cells expressing both Sox2 and Sox17 at very low levels (Sox2-Sox17-low cells) was observed (Figure 6F, red dots). These cells are likely unspecified cells or were not specified to either fate; these unspecified cells are eliminated through cell competition. Importantly, similar Sox2-Sox17-low cells were also observed in wild-type cells of Z-VAD-treated wild-type ⇔ *Tead1*^−/−^ embryos (Figure 6E), suggesting that unspecified cells are also produced during the normal developmental process and that such cells are eliminated via cell competition.

**Figure 6.**
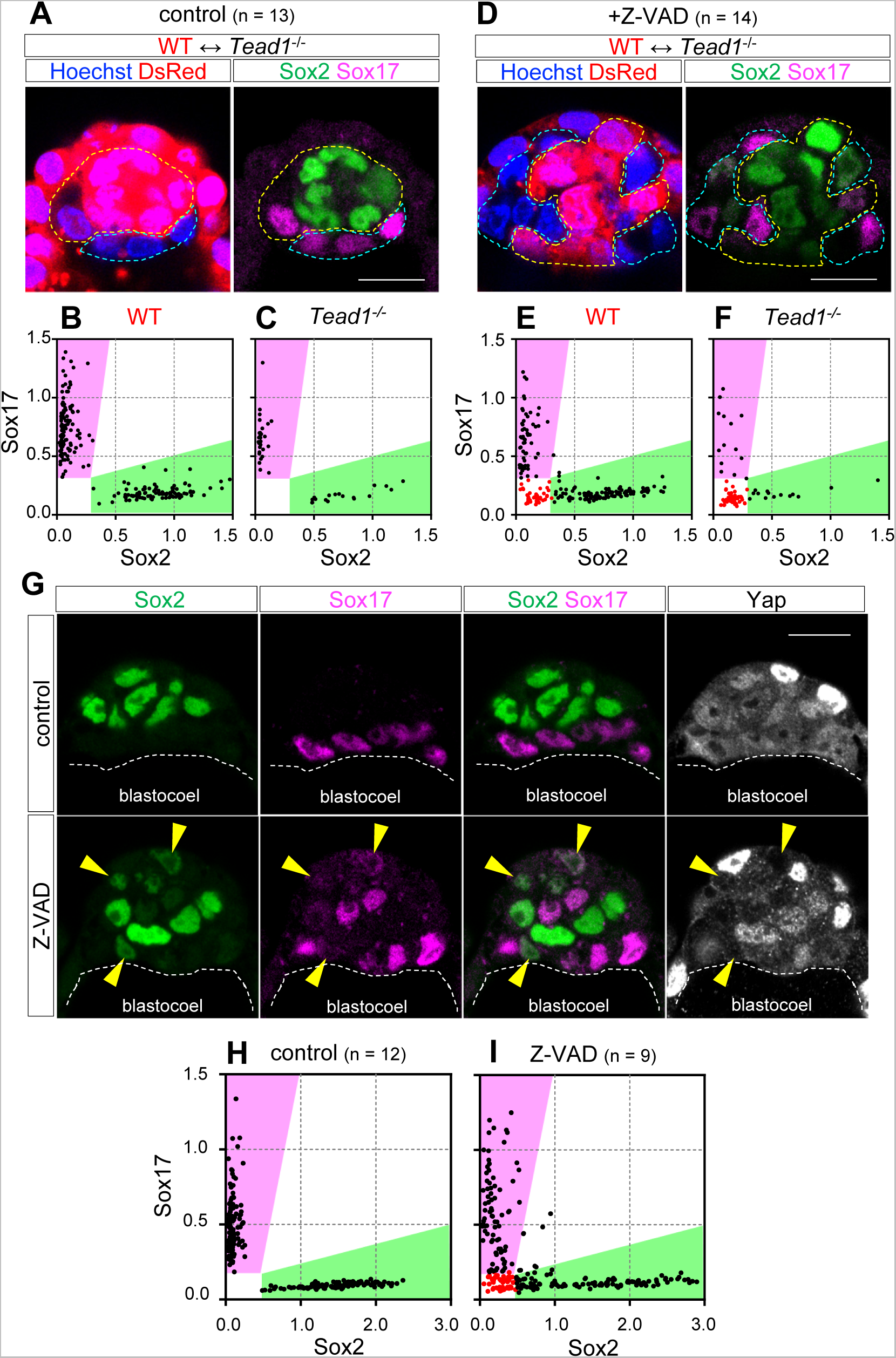
Cell competition eliminates unspecified cells from epiblast. (A-F) Expression of Sox2 and Sox17 in control (A-C) and Z-VAD-treated (D-F) wild-type ⇔ *Tead1*^−/−^ embryo. Wild-type and *Tead1*^−/−^ cells are indicated with yellow and cyan dashed lines, respectively. Quantification of Sox2 and Sox17 signals in wild-type (B, E) and *Tead1*^−/−^ (C, F) cells. (G) Z-VAD-treated wild-type embryo showing disrupted organization of epiblast and primitive endoderm. Quantification of Sox2 and Sox17 signals in control (H) and Z-VAD-treated (I) embryos. Arrowheads indicate weak double-positive cells showing weak nuclear Yap. Scale bars are 20 µm for panels A, D, G.

To further evaluate this hypothesis, we next treated normal embryos with Z-VAD. In control embryos, all cells were single-positive for either Sox2 or Sox17, and these cells were segregated to the inner and outer (facing the blastocoel) positions, respectively, indicating proper cell fate specification and tissue organization (Figure 6G, H). In Z-VAD-treated embryos, Sox2-Sox17-low cells were also observed, supporting that unspecified cells are produced during normal development and that these cells are eliminated via cell competition (Figure 6G, I). The ICM sizes of Z-VAD-treated embryos tended to be larger than those of normal embryos (Figure 6G). A scattered presence of weak Sox2-Sox17 double-positive cells (arrowheads in Figure 6G), which also showed weak nuclear Yap, in the inner cells perturbed formation of the spatially segregated and properly organized epiblast and primitive endoderm. Thus, elimination of unspecified cells via cell competition is important for proper organization of the epiblast and primitive endoderm.

## Discussion

### Epiblast formation is achieved by Tead-Yap-dependent expression of pluripotency factors and elimination of unspecified cells via cell competition

In this study, we found that Hippo signaling regulates naïve pluripotency in preimplantation embryos. We also determined a quality control mechanism via cell competition occurring during formation of the epiblast. These findings are summarized in the model shown in Figure 7. Up to the early blastocyst stage, Hippo signaling regulates the first cell fate specification (Sasaki, 2017). In inner cells, active Hippo signaling inhibits nuclear localization of Yap and inactivation of Tead promotes the ICM fate (Hirate et al., 2012; Nishioka et al., 2009). After specification of the ICM, the Hippo signal is gradually attenuated in the ICM, allowing nuclear accumulation of Yap. Thus, from the mid-blastocyst stage, a gradual increase in Tead activity promotes the expression of pluripotency factors. At the mid-blastocyst stage, the levels of nuclear Yap (i.e. Tead activity) in ICM cells are highly variable and correlate with the expression levels of pluripotency factors, indicating that induction by Tead-Yap is a major mechanism of pluripotency factor expression at this stage. The strong correlation between nuclear Yap and Sox2 expression likely reflects direct regulation of *Sox2* by Tead-Yap, as observed in ES cells (Lian et al., 2010). Development of *Tead1*^−/−^ or *Yap1^−/−^* embryos up to post-implantation stage does not imply that Tead-Yap is dispensable for epiblast formation. Presence of three other *Tead* genes or *Wwtr1* likely partially compensates for the absence of *Tead1* or *Yap1*, respectively, and supports production of smaller number of epiblast cells. Because of the lethality of the embryos lacking all four *Tead* genes or both *Yap1* and *Wwtr1* before the late blastocyst stage (Nishioka et al., 2009; H. S. unpublished observation), it remains unknown whether they are essential or not. Tead-Yap-dependent expression of pluripotency factors is an evolutionarily conserved mechanism, as human blastocysts also show nuclear localization of YAP in the ICM (Qin et al., 2016).

**Figure 7.**
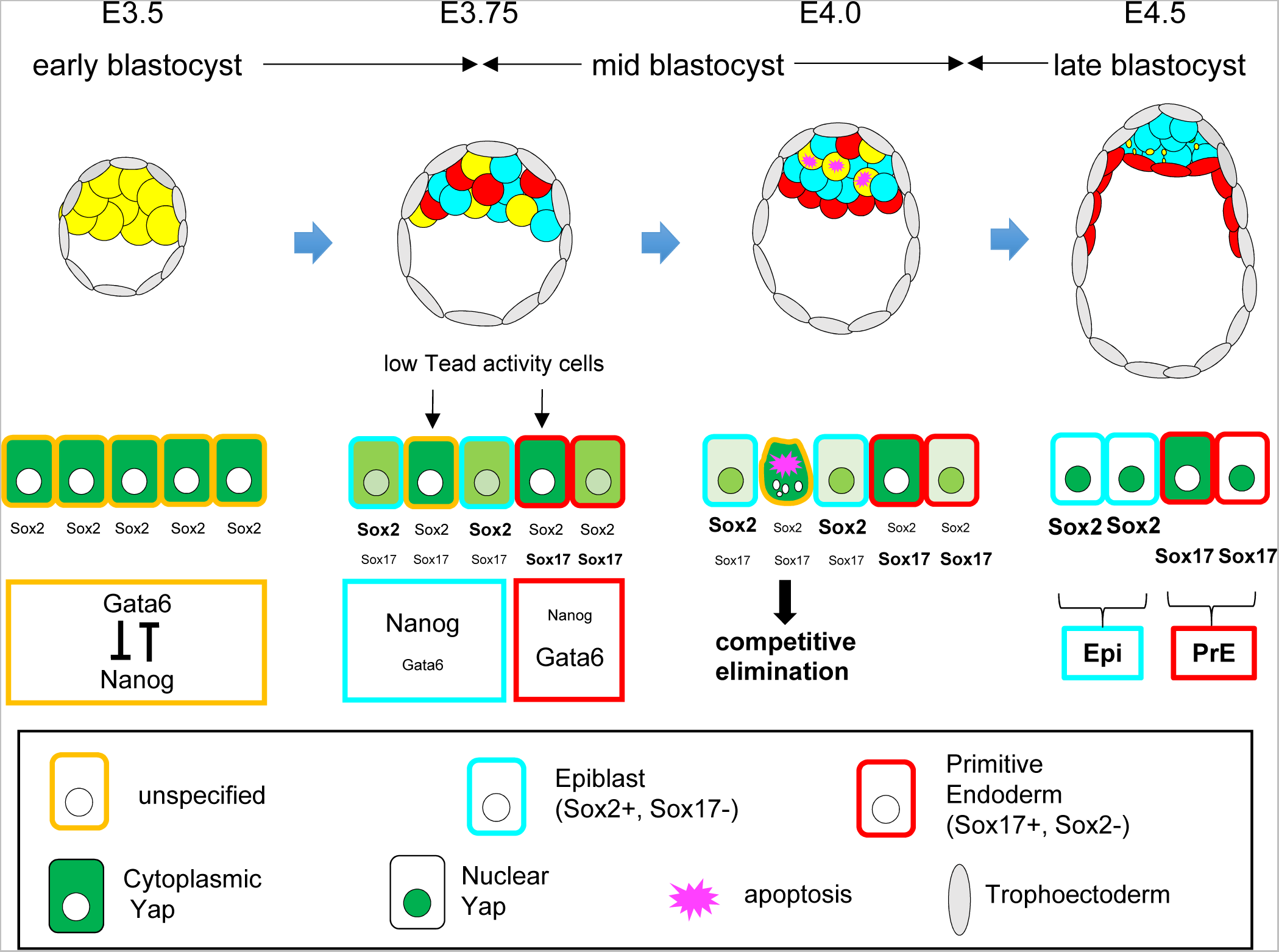
Model of the roles of Hippo signaling and cell competition during epiblast formation in preimplantation mouse embryos. Top panels show spatial distribution of ICM/epiblast/primitive endoderm cells. Middle panels show gene expression/differentiation status of ICM cells. The sizes of characters indicate expression levels. Green color indicates subcellular distribution of Yap. Color codes of the frames of the cells indicate cell differentiation status. See details for discussion.

Variations in Tead activities cause two distinct responses among ICM cells (Figure 7). By the mid-blastocyst stage, expression of Nanog and Gata6, initially co-expressed in the ICM, are segregated into a salt-and-pepper pattern through mutual exclusion mechanisms (Chazaud and Yamanaka, 2016). If low Tead-activity cells, which express low level of pluripotency factors, express high levels of the primitive endoderm regulators (e.g. Gata6, Sox17, etc.), then these cells become primitive endoderm. Among primitive endoderm-specified cells, differences in Tead activities do not trigger cell competition. In contrast, if low Tead-activity cells also fail to express primitive endoderm regulators, then such unspecified cells are recognized as low fitness epiblast cells by neighboring high Tead-activity epiblast cells and are eliminated via cell competition. It is important to note that cell competition occurs only among epiblast cells. Expression of pluripotency factors, or most likely pluripotency, and Myc, which are regulated by Tead activity, appear to be important factors that cooperatively control cell competition. At the late blastocyst stage, the remaining cells which express high level of pluripotency factors acquire naïve pluripotency, likely by activating the robust transcription factor network for naïve pluripotency (Dunn et al., 2014; Niwa, 2018). Because this network does not contain Tead-Yap, expression of pluripotency factors becomes less sensitive to variations in Tead activity. Indeed, expression of the pluripotency factors Sox2 and Oct3/4 became less variable at the late blastocyst stage, while Yap and Nanog maintained the high level of variation (Figure 4H, I, Figure S3G, H).

How is Yap regulated in the forming epiblast? Dynamic cellular behaviors likely contribute to variations in nuclear Yap levels. Division of a cell should cause dynamic changes in the morphology of and/or mechanical forces applied to dividing and neighboring cells. These changes affect the cytoskeletal organization and/or nuclear shape of the cells, both of which control nuclear entry of Yap through both Hippo signaling-dependent and -independent mechanisms (Dupont et al., 2011; Elosegui-Artola et al., 2017; Wada et al., 2011; Zhao et al., 2012). Attenuation of Hippo signaling by reduced expression of the key Hippo pathway components *Amot* and *Lats2* in the epiblast (this work and Boroviak et al., 2015) is a likely mechanism responsible for the gradual increase in nuclear Yap. Expression of *Amot* and *Lats2* are also weak in ES cells (Galonska et al., 2015), suggesting that the acquisition of pluripotency attenuates Hippo signaling to maintain pluripotency. Changes in cellular morphology may also contribute to the gradual increase in nuclear Yap, as the nuclear shapes of the ICM/epiblast cells change from spherical to flat from the early to late blastocyst stages. Increased hydrostatic pressure in the blastocyst cavity may increase tension on the forming epiblast cells.

### Cell competition controls epiblast quality in two steps

Our study revealed that cell competition eliminate unspecified cells during the formation of the naïve epiblast, while previous studies showed that cell competition controls epiblast quality at the post-implantation stage at approximately E6.0 (Claveria et al., 2013; Sancho et al., 2013) during the exit from formative pluripotency (Smith, 2017). Therefore, epiblast quality is controlled via cell competition at two stages: during establishment of naïve pluripotency and exit from formative pluripotency. Why are these two quality-control steps required? In the first quality-control step, likely because of the nature of preimplantation development, cell differentiation from the ICM to the epiblast and primitive endoderm is highly dynamic: gene expression is heterogeneous and dynamically fluctuates, and the timing of cell differentiation is variable (Ohnishi et al., 2014; Saiz et al., 2016; Xenopoulos et al., 2015). Strong variations in Tead activity result in the production of large numbers of unspecified cells. Cell competition is required to eliminate such unnecessary cells produced by developmental fluctuations to establish a naïve epiblast. For the second quality-control step, as described previously, cell competition is required to eliminate the prematurely differentiation-primed cells (Diaz-Diaz et al., 2017) and defective cells produced by the increased proliferation rate or karyotypically abnormal cells to prevent these cells from contributing to the germline (Bowling et al., 2018). Thus, two quality-control steps via cell competition may function as safeguards against developmental fluctuations and errors to ensure the production and expansion of a high-quality epiblast to support correct embryonic and germ line development. Although *Bok^−/−^Bax^−/−^Bak^−/−^* triple-knockout mice, which lack intrinsic apoptosis ability, formed a morphologically normal ICM and post-implantation epiblast (Ke et al., 2018), it remains unclear whether the qualities of the epiblasts are also normal, as many of the mutants exhibited various morphological abnormalities as well as severe growth retardation and death at E10.5 (Ke et al., 2018).

### Pluripotency and Myc regulate cell competition

Although differences in Tead activity triggered cell competition in the epiblast, it appeared that expression of pluripotency factors, or most likely pluripotency, and Myc, but not Tead activity itself, control cell competition. The importance of pluripotency is also suggested by the occurrence of cell competition only in the epiblast. Similarly, elimination of aneuploidy cells occurred only in the epiblast (Bolton et al., 2016). Thus, the mechanism of cell competition differs from the cell competition regulated by Tead-Yap in *Drosophila* and NIH3T3 cells, in which Tead activity and downstream Myc cooperatively control cell competition (Mamada et al., 2015; Neto-Silva et al., 2010; Ziosi et al., 2010). Pluripotency-regulated cell competition resembles cell competition in ES cells and post-implantation epiblasts. In ES cells, the pluripotency status determines Myc expression levels, and Myc controls cell competition (Diaz-Diaz et al., 2017). In that study, however, it was not clearly demonstrated whether Myc alone regulates cell competition downstream of pluripotency, or if Myc and pluripotency cooperatively regulate competition. In preimplantation epiblasts, Myc is involved in but not the only factor controlling cell competition downstream of Tead activity. We predict that pluripotency and Myc cooperatively regulate competition. Additional studies are needed to test this hypothesis. Because all cell competitions in pre- and post-implantation epiblasts and in ES cells appear to compare pluripotency status, it is important to clarify the similarities and differences among these cell competition events, as well as determine the mechanisms by which the cells recognize the pluripotent status of their neighbors.

## Acknowledgments

We thank Dr. Hitoshi Niwa for critical reading of the manuscript, Makoto Adachi for anti-Amotl2 antibody, Ken-ichi Yagami and RIKEN BRC for R26GRR mouse, and Feng Zhang and Addgene for plasmids. This work was supported by JSPS KAKENHI (Grant Numbers JP15H01495, JP17H03682, JP17H05618, JP23247036, JP26112715), research grants from the Mitsubishi Foundation, Uehara Memorial Foundation, and Takeda Science Foundation to H.S. and JSPS KAKENHI Grant Number JP16K18978 to M.H.

## Author contributions

Conceptualization, M.H. and H.S.; Methodology, M.H. and H.S.; Investigation, M.H.; Visualization, M.H.; Writing – Original Draft, M.H. and H.S.; Writing – Review & Editing, M.H. and H.S.; Funding Acquisition, M.H. and H.S.

## STAR Methods

### CONTACT FOR REAGENT AND RESOURCE SHARING

Further information and requests for resources and reagents should be directed to and will be fulfilled by the Lead Contact, Hiroshi Sasaki.

### EXPERIMENTAL MODEL AND SUBJECT DETAILS

#### Mice

Mice were housed in environmentally controlled rooms in the Animal Facility of the Frontier of Biosciences, Osaka University. All experiments with mice and recombinant DNA were performed with approval from the Animal Care and Use Committee of the Graduate School of Frontier Biosciences, Osaka University and Gene Modification Experiments Safety Committee of Osaka University, respectively.

We used 3-week-old B6D2F1/Slc, (C57BL/6NCrSlc x DBA/2CrSlc)F1, female mice purchased from Japan SLC (Hamamatsu, Japan) or colonized in our facility to obtain embryos. B6D2F1/Slc female mice were superovulated by intraperitoneal injection of 10 IU pregnant mare serum gonadotropin and human chorionic gonadotropin at 48-h intervals and crossed with B6D2F1/Slc male mice to obtain wild-type embryos, or homozygotic R26GRR (C57BL/6N) (Hasegawa et al., 2013) or R26RR (mixed background of C57BL/6N × Slc:ICR) male mice to perform mosaic analyses. R26RR mice were generated by removing the GFP cassette from R26GRR mice by introducing NLS-Cre mRNA into R26GRR 1-cell stage zygote by electroporation (see below).

### METHOD DETAILS

#### Embryo culture and manipulation

One-cell-stage zygotes were collected in M2 medium from the oviduct ampulla at the day of plug (E0.5) and treated with hyaluronidase to remove the cumulus cells. Two-cell-stage embryos were collected from the E1.5 oviduct by M2 medium flashing. Eight-cell or later stage preimplantation embryos were collected by M2 medium flashing of the uterine. All embryos were cultured in 10-µL drops of KSOM covered with 5 µL mineral oil in each well of Nunc™ MiniTrays with Nunclon™ Delta surface at 37°C in a 5% CO_2_ incubator.

#### NLS-Cre mRNA preparation

cDNA coding for the nuclear localization signal (RKKKRKV; CCCAAGAAGAAGAGGAAGGTG) tagged with Cre recombinase was cloned into the pCS2 expression vector (pCS2-NLS-Cre). NLS-Cre mRNA was synthesized by *in vitro* transcription from the pCS2-NLS-Cre vector by using the mMESSAGE mMACHINE SP6 Transcription Kit (Thermo Fisher Scientific, Waltham, MA, USA) following the manufacture’s protocols. The mRNA was purified by phenol-chloroform-isoamyl alcohol extraction and isopropanol precipitation, and resolved in Opti-MEM I (Gibco, Grand Island, NY, USA).

#### sgRNA design and synthesis

The sgRNAs for each gene were designed by using GPP Web Portal developed by the Broad Institute and/or mm10 CRISPR targets database (Sunagawa et al., 2016). Each target sequence was 21–40 nucleotides of 70 nucleotide-long forward primers (see below). We prepared PCR products amplified from the pX330-U6-Chimeric_BB-CBh-hSpCas9 vector (Cong et al., 2013) by using a 70 nucleotide-long forward primer (Eurofins Genomics, Table S1) containing a T7site and sgRNA specific and common sequence and 26 nucleotide-long Common gRNA Reverse primer (Table S1) with KOD Fx neo DNA polymerase (Toyobo, Osaka, Japan). The thermal cycler program was 96°C for 1 min, followed by 96°C for 10 s, 63°C for 10 s, and 68°C for 30 s for 40cycles, and then 68°C for 1 min, with the final temperature held at 4°C. The PCR amplicon was purified by ethanol precipitation and rinsed in 70% ethanol. sgRNAs were directly synthesized from the purified PCR amplicon using the MEGAshortscript T7 Transcription Kit (Thermo Fisher Scientific) following the manufacture’s protocol. sgRNAs were purified by phenol-chloroform-isoamyl alcohol extraction and isopropanol precipitation, and resolved in RNase-free water.

#### Zygote electroporation

To knockout genes at the 1-cell stage, we performed zygote electroporation using the CRISPR/Cas9 system method developed in our previous study (Hashimoto et al., 2016). The zygotes were aligned in the 1-mm gap of an LF501PT1-10 electrode (BEX Co., Ltd., Tokyo, Japan) filled with freshly prepared 5 µL of 200 ng/µL Alt-R^®^ S.p. Cas9 Nuclease V3 and 100 ng/µL sgRNA in Opti-MEM. The electric conditions were to 2 sets of 5 pluses of 3 ms at 25 V, with inverting polarity between sets by using the CUY21 EDIT II electroporator (BEX Co., Ltd., Tokyo, Japan). For Cre-loxP recombination to remove the GFP cassette from the R26GRR embryo to generate R26RR mice, the R26GRR zygotes were placed in a solution of 25 ng/µL NLS-Cre mRNA in Opti-MEM I and electroporated by 2 sets of 3 pluses of 3 ms at 25 V, with inverting polarity between sets.

#### Microinjection

A freshly prepared solution of 200 ng/µL Alt-R^®^ S.p. Cas9 Nuclease V3 and 100 ng/µL each sgRNA in Opti-MEM were injected into the cytoplasm of 2-cell stage embryos at E1.5. NLS-Cre mRNA was used at a final concentration of 1 ng/µL for the R26GRR embryos. The embryos were incubated in M2 medium during microinjection at room temperature. The maximum number of injected embryos in a single experiment was 50, and each experiment was conducted in less than 20 min.

#### Inhibitor treatments

To force the expression of pluripotent markers, the embryos were cultured in KSOM containing 2i (1.0 µM MEK inhibitor PD0325901 and 3.0 µM Gsk3 beta inhibitor CHIR99021) and 0.04% dimethyl sulfoxide (DMSO) from the 8-cell stage until the late blastocyst stage. For Myc inhibition, embryos were cultured in KSOM containing 55 µM of c-Myc inhibitor and 0.11% DMSO from the mid-blastocyst stage to the late blastocyst stage. To inhibit apoptotic cell death, embryos were cultured the KSOM containing 200 µM of Z-VAD-FMK and 0.4% DMSO from the early blastocyst stage to the late blastocyst stage. All control embryos were cultured in KSOM containing the same concentration of DMSO for the same duration for each experiment.

#### Immunofluorescent staining and confocal image acquisition

All antibodies used in this study are listed in the Key Resources Table. Immunofluorescent staining of embryos was performed following standard protocols. Embryos were fixed in 4% paraformaldehyde in phosphate-buffered saline (PBS) for 5 min at room temperature, and then washed and permeabilized with 0.1% Triton X-100 in PBS (0.1% PBT) for 1 min twice at room temperature, blocked with 2% donkey serum in 0.1% PBT (blocking solution), and incubated overnight with primary antibodies diluted at 1:100 in blocking solution at room temperature. After washing in 0.1% PBT for 1 min three times, the embryos were incubated with the secondary antibodies, and then Hoechst for nuclear staining at a 1:1000 dilution in 0.1% PBT for more than 1 h at room temperature. Immunofluorescence-stained embryos were placed in a PBS drop on a glass base dish and confocal images were obtained by using a Nikon A1 inverted confocal microscope (Tokyo, Japan) and analyzed by using NIS-Elements AR analysis (Nikon), IMARIS (Bitplane, Belfast, UK), or ImageJ software (NIH, Bethesda, MD, USA).

#### Alexa Fluor^®^ labeling of primary antibodies

For simultaneous immunostaining of multiple antibodies raised from the same host animals, we labeled the primary antibodies with Alexa Fluor^®^ in advance by using Molecular Probes^®^ Antibody Labeling Kits (Eugene, OR, USA) according to the manufacturer’s protocols. In this study, Sox2-Alexa647, Sox17-Alexa488 (both are raised in goat) and Yap1-Alexa488, cleaved Caspase3-Alexa647, Nanog-Alexa568 (all are raised in rabbit) were labeled in advance.

### QUANTIFICATION AND STATISTICAL ANALYSIS

#### Quantification of cell competition

The ratio of DsRed-positive or DsRed-negative Sox2-expressing cells in mosaic embryos was estimated by manually counting the number of each cell in the embryos by NIS-Elements AR analysis.

#### Quantification of confocal images

The obtained confocal images were quantitatively analyzed with IMARIS software. To quantify the nuclear localizing Yap, Sox2, Nanog, Oct3/4, Sox17, and active-Yap in the ICM/Epiblast, the signal intensities of each marker were divided by the Hoechst signal intensity to normalize the signal attenuation on the Z-axis. The expression levels of each marker were the means of signal intensities in the regions of Sox2-positive 6-µm spheres in diameter quantitated by using a spots program. To quantify Amotl2, Amot, and p-Yap signals, the cells of Sox2-positive cells were manually cropped by using ImageJ software from a single 2D confocal image, and the means of signal intensities in the cropped areas were calculated using ImageJ software.

#### Statistical analysis

All statistical analyses in this study were performed using GraphPad Prism6 software (GraphPad, Inc., La Jolla, CA, USA). The number of cells or embryos analyzed (n), presented error bars (SD or SEM), statistical analyses, and *p* values are all stated in each figure or figure legend.

---

**Figure S1. Related to Figure1.**
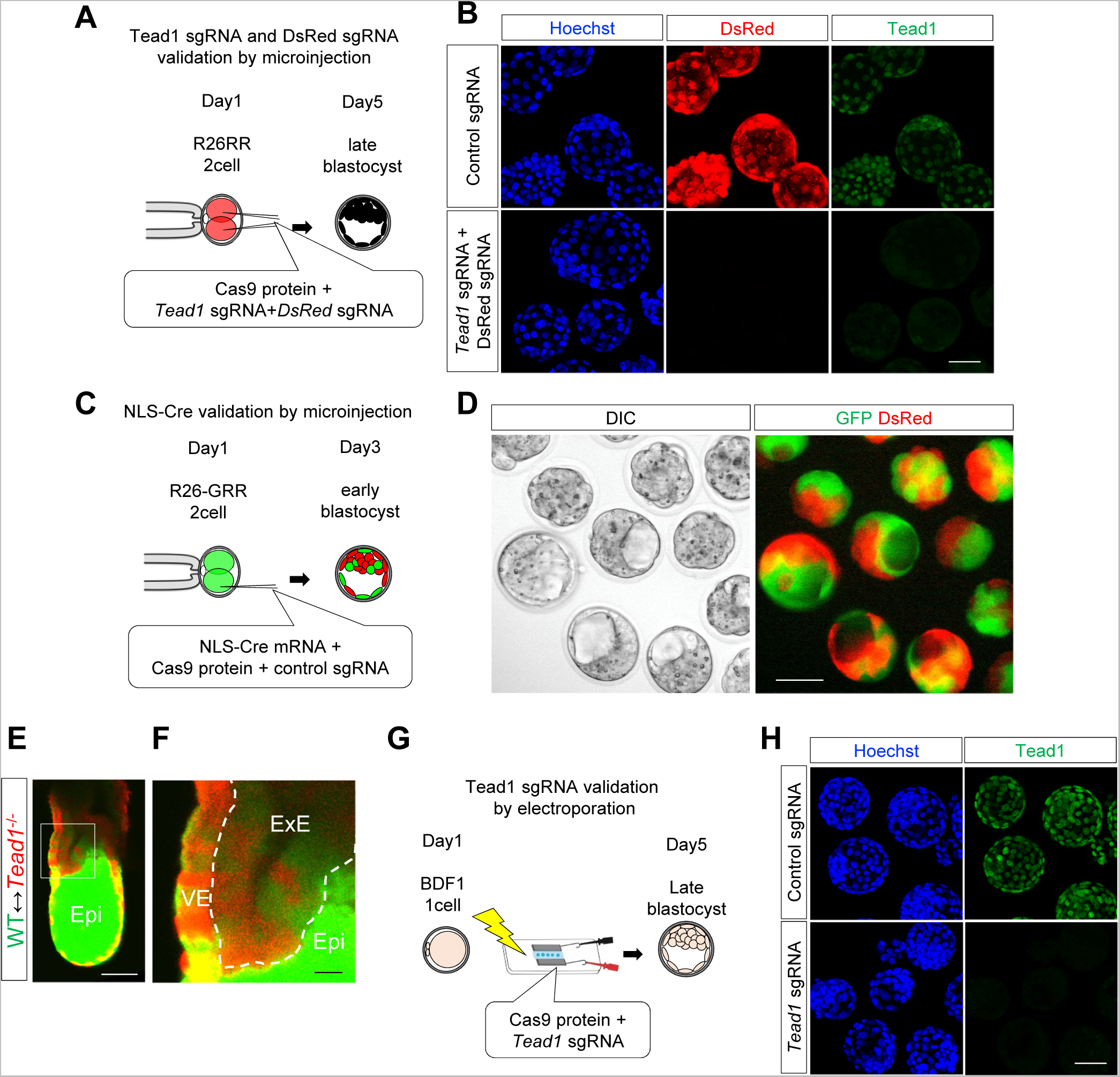
Efficiencies of genome editing and Cre recombinase. (A and B) Efficiency of genome editing for *Tead1* and *DsRed* by microinjection at the 2-cell stage. (A) Scheme of experiments. (B) Representative embryos showing absence of DsRed fluorescence and strong reduction in Tead1 protein levels in all cells. Strong reduction in Tead1 signals indicates disruption of *Tead1,* and very faint signals likely reflect remaining Tead1 proteins produced before genome editing at the 2-cell stage. (C and D) Efficiency of Cre recombinase by microinjection of mRNA at the 2-cell stage. (C) Scheme of experiments. (D) Representative embryos showing color conversion of the Cre reporter mouse, R26-GRR, in all embryos. (E and F) Contribution of *Tead1*^−/−^ cells to extra-embryonic ectoderm in wild-type ⇔ *Tead1*^−/−^ embryos. (E) Enhanced image of the wild-type ⇔ *Tead1*^−/−^ embryo shown in Figure 1C. (F) Enlargement of the boxed area of (E), showing contribution of *Tead1*^−/−^ (red) cells to extra embryonic ectoderm (ExE). VE: visceral endoderm, Epi: epiblast. (G, H) Efficiency of genome editing for *Tead1* by electroporation at the 1-cell stage. (G) Scheme of experiments. (H) Representative embryos showing absence of Tead1 proteins in all cells. Scale bars represent 100 µm for panel E, 50 µm for B, D, H, and 20 µm for F.

**Figure S2. Related to Figure 2.**
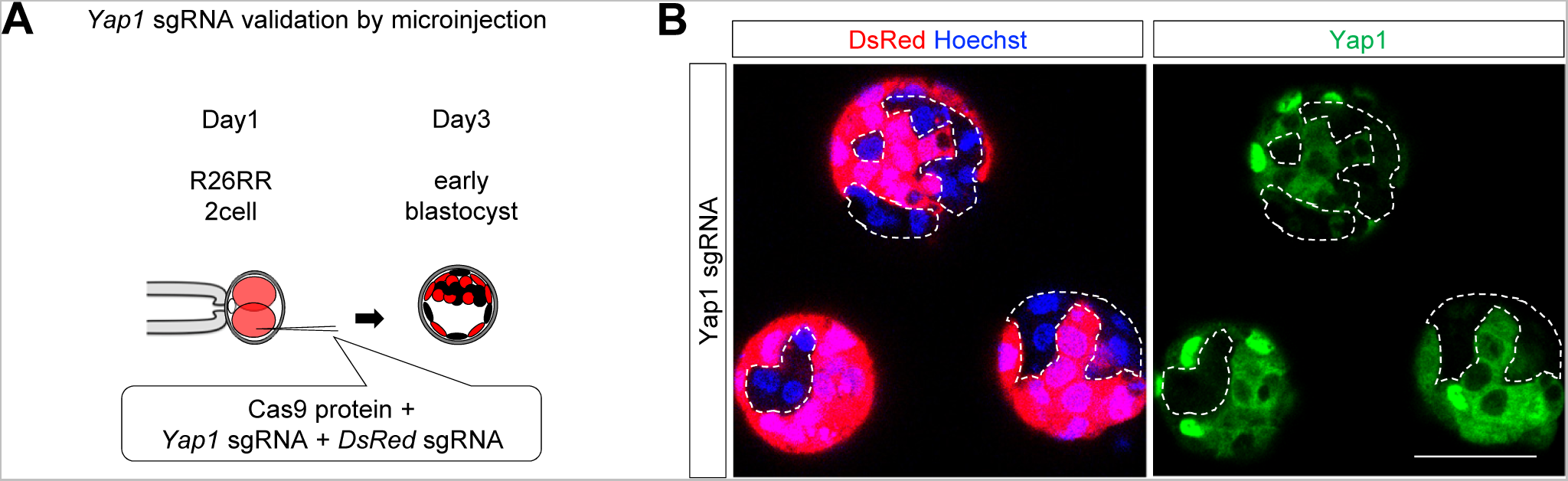
Efficiencies of genome editing for Yap1. (A) Scheme of experiments. (B) Representative embryos showing strong reduction in Yap1 protein levels in all manipulated cells lacking DsRed signal. Strong reduction in Yap1 signals indicates disruption of *Yap1*, and very faint signals likely reflect remaining Yap1 proteins produced before genome editing at the 2-cell stage. Scale bars represent 50 µm.

**Figure S3. Related to Figure 4.**
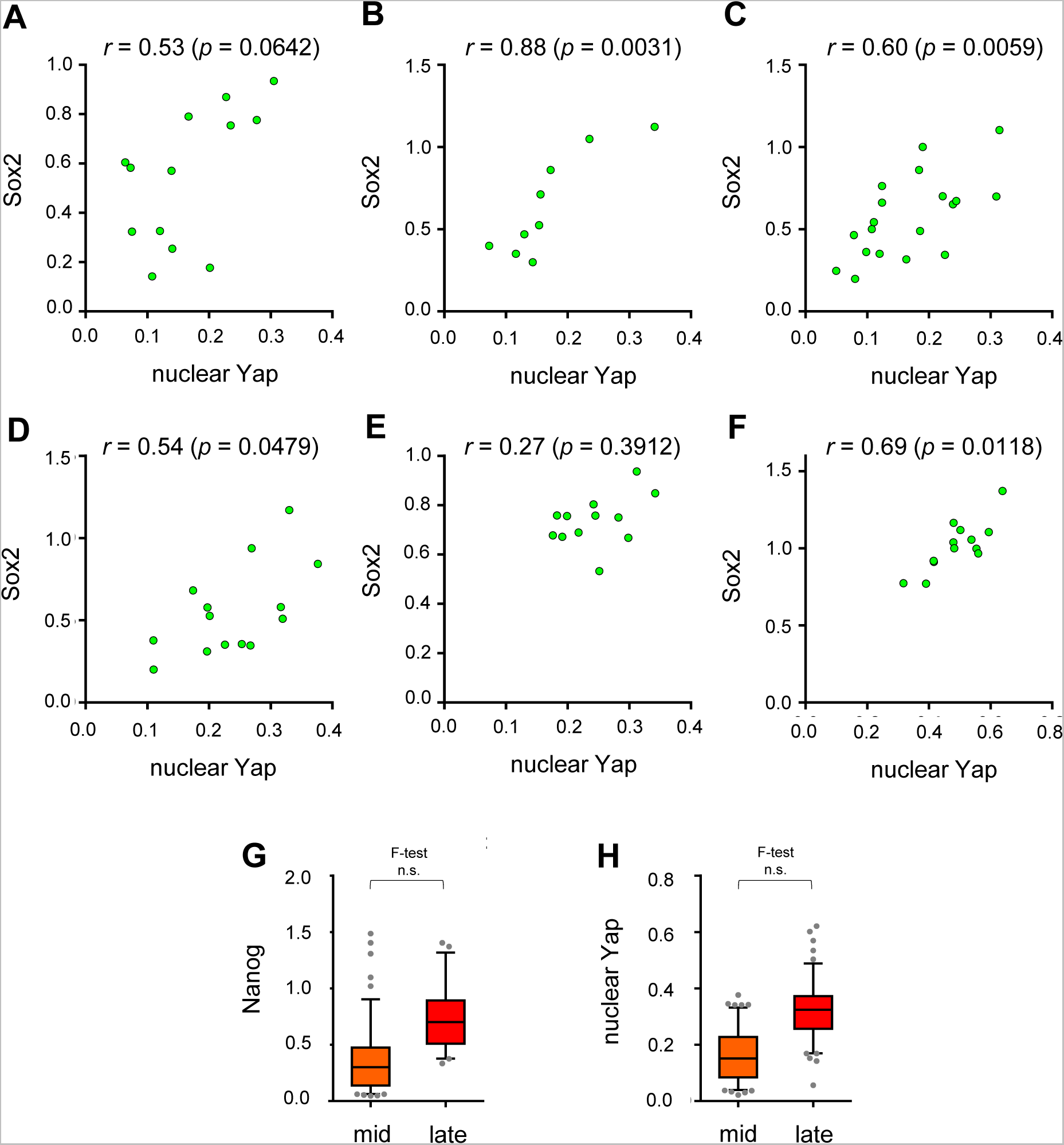
Sox2 and nuclear Yap signals in the ICM are highly variable in individual embryos at the mid blastocyst stage. (A–F) Graphs showing distribution of Sox2 and nuclear Yap signals in the ICM of an embryo. Each graph represents the data from one embryo. (G, H) Comparison of variations of signal intensities between mid- and late blastocyst stages. Distribution of signal intensities for Nanog (G), and nuclear Yap (H) at mid- and late blastocyst stages. Data are shown as box and whisker plots. Box and the horizontal line indicate the range of central 50% and median, respectively. Whiskers represent the range from 5% to 95%. n.s.: not significant, *F*-test.

**Figure S4. Related to Figure 5.**
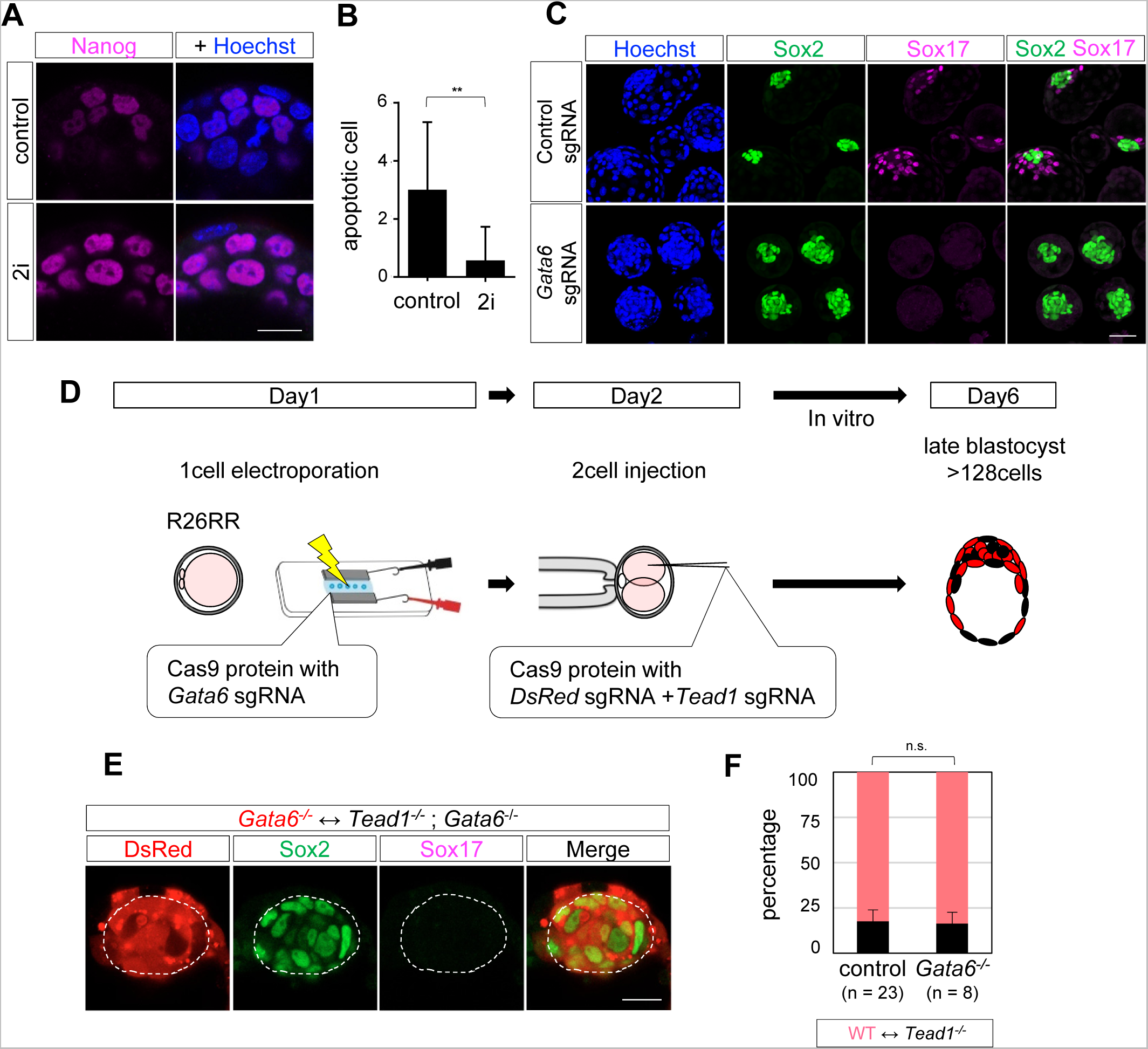
Pluripotency regulates cell competition. (A) Effects of 2i-treatment on expression of Nanog. (B) Quantification of number of cleaved caspase 3-positive ICM cells. The number of cleaved caspase 3-positive cells were reduced in 2i-treated wild-type embryos. Data are represented as the mean ± SD. *\*\*p<* 0.01, Mann-Whitney *U*test. (C) Representative *Gata6^−/−^* embryos produced by genome editing at the 1-cell stage, showing absence of Sox17-positive primitive endoderm cells and increase in the number of Sox2-positive epiblast cells. (D) Scheme of the experiments used for *Gata6^−/−^* ⇔ *Tead1*^−/−^;*Gata6^−/−^* mosaic embryos. (E) Representative *Gata6^−/−^* ⇔ *Tead1*^−/−^;*Gata6^−/−^* embryos showing absence of *Tead1*^−/−^ cells in the epiblast. (F) Quantification of the percentage of *Tead1*^−/−^ cells in *Gata6^−/−^* ⇔ *Tead1^−/−^;Gata6^−/−^* embryos. *Gata6*^−/−^ mutation did not rescue elimination *of Tead1^−/−^* cells. Data are represented as the mean ± SEM. Mann-Whitney *U*test Scale bars represent 50 µm for panel C, and 20 µm for A, E.

**Figure S5. Related to Figure 5.**
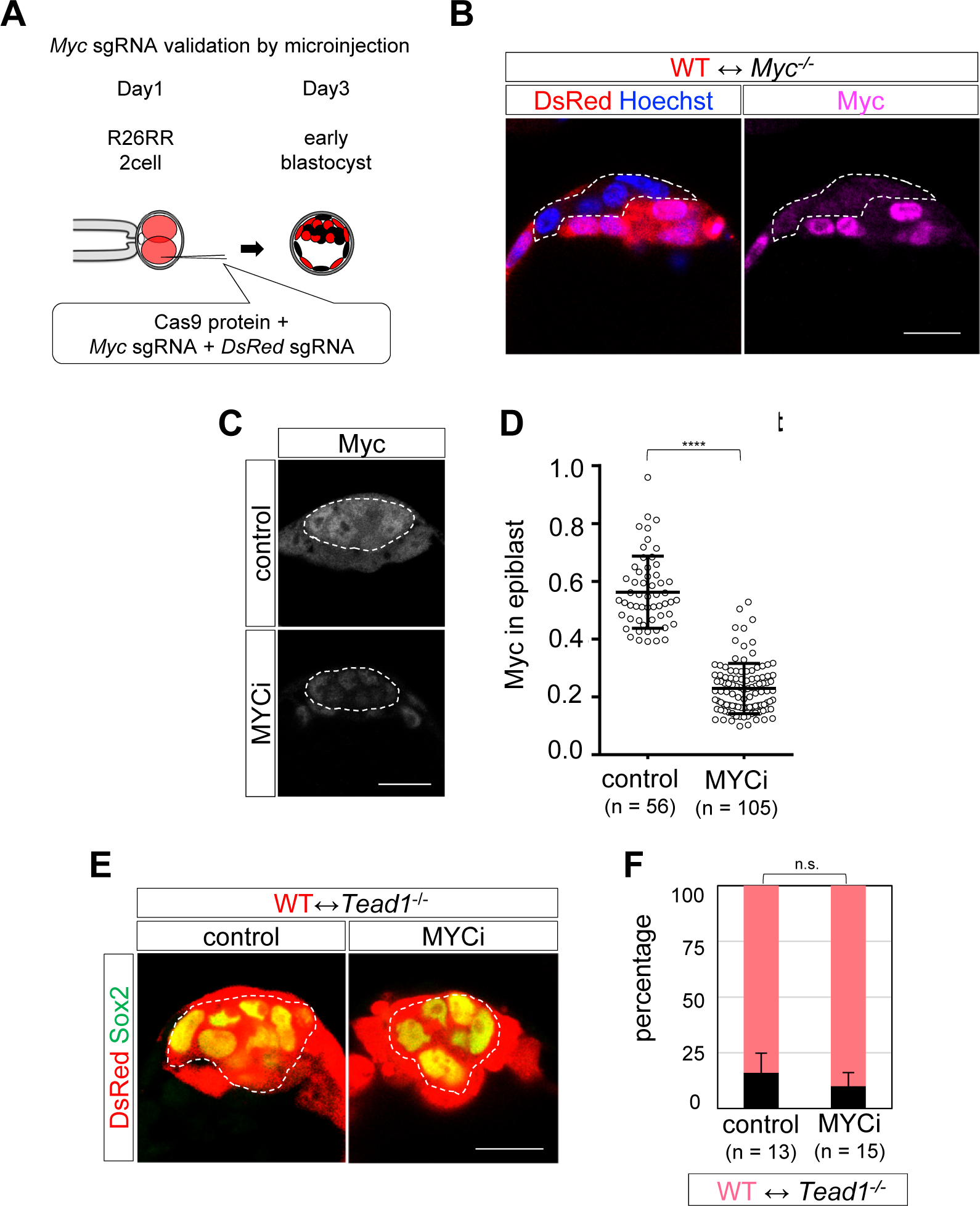
Contribution of Myc to cell competition. (A and B) Efficiency of genome editing for *Myc* by microinjection at the 2-cell stage. (A) Scheme of experiments. (B) Representative embryo showing clear reduction of Myc protein levels in DsRed-negative injected cells. Weak signals likely reflect background level signals of the anti-Myc antibody. (C) MYCi-treated embryo showing reduced expression of Myc. (D) Quantification of the Myc signal intensities in epiblast cells. (E) MYCi-treated wild-type ⇔ *Tead1*^−/−^ embryo showing loss of *Tead1*^−/−^ cells from the epiblast. (F) Quantification of *Tead1*^−/−^ cells in epiblast. Data are represented as mean ± SD (D) or SEM (F). Students’ *t*-test (D), and Mann-Whitney *U*test (F). **** *p* < 0.0001 Scale bars represent 20 µm for B, C, E.

**Table S1.**
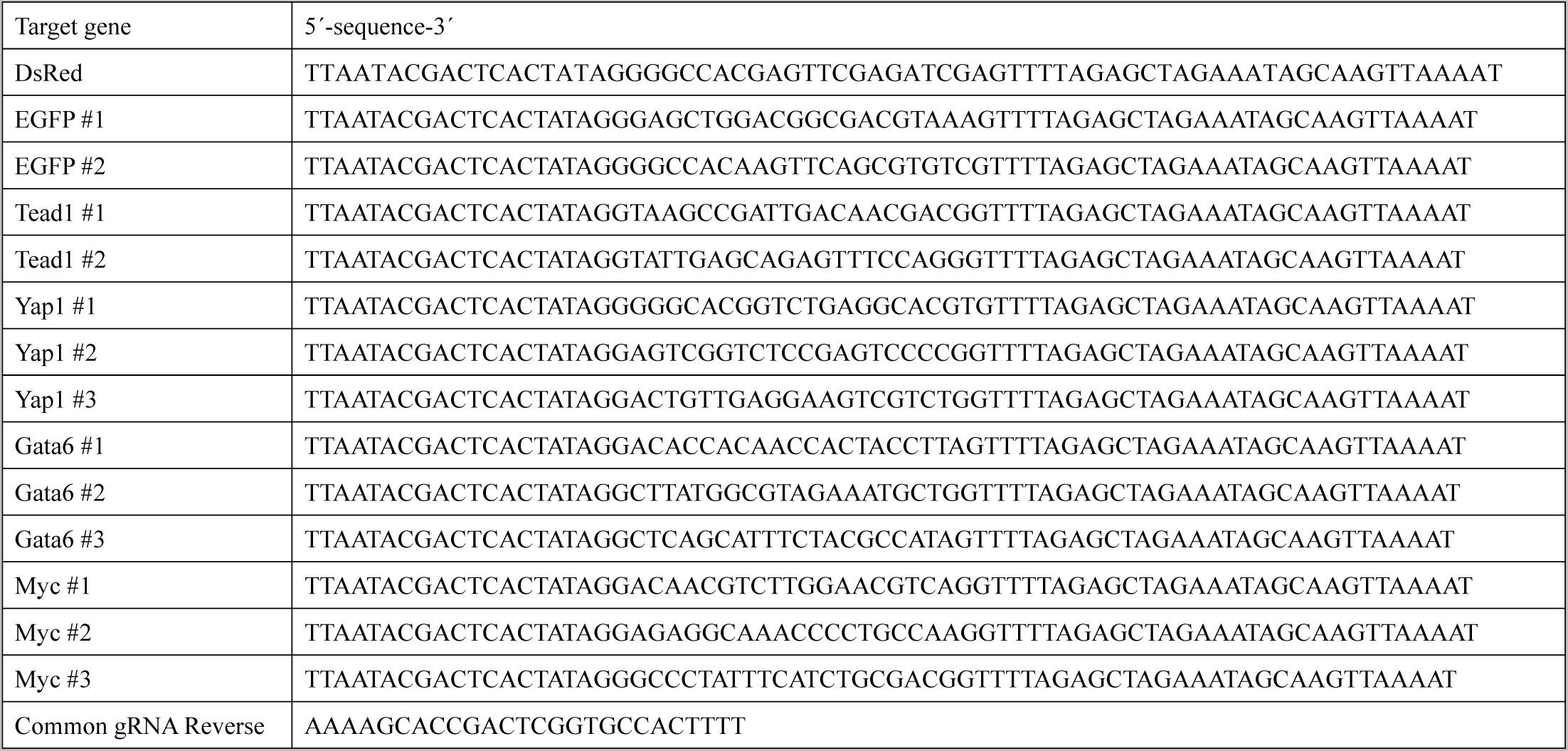
Related to KEY RESOURCES TABLE and STAR Methods. The list of oligonucleotides used to amplify the gRNA templates by PCR

